# A population genetic study of *Fusarium graminearum*, the causal agent of Fusarium Head Blight, provides insights into patterns of inoculum dispersal in wheat fields at a regional scale

**DOI:** 10.1101/2025.07.11.664332

**Authors:** Toan Bao Hung Nguyen, Gaétan Le Floch, Amandine Henri-Sanvoisin, Sylvie Treguer, Marie Foulongne-Oriol, Adeline Picot

## Abstract

*Fusarium graminearum* is the primary pathogen responsible for Fusarium Head Blight (FHB) in cereals, causing yield losses and mycotoxin contamination. The present study investigated the genetic diversity and population structure of *F. graminearum* from wheat grains and maize residues, which were the primary inoculum sources of FHB. A total of 588 isolates from four different fields were genotyped using a sequence-based microsatellite genotyping method of 34 SSR markers (SSRseq) and characterized for chemotype distribution. The results revealed high genetic diversity with 453 distinct multilocus genotypes (MLGs) without significant genetic differentiation by geographic location or substrate, suggesting a single metapopulation at the regional scale. This absence of substrate genetic differentiation strongly supports the fact that residue-borne inoculum contributed to FHB infection, although we failed to demonstrate any field-specific dispersal pattern from residues to grains. The findings underscore the importance of maize residues as a critical inoculum source and highlight the need to consider both local and regional sources in FHB management.

## Introduction

*Fusarium graminearum* sensu stricto (teleomorph: *Gibberella zeae*) is considered to be the major harmful pathogen responsible for Fusarium Head Blight (FHB), an important disease affecting several cereals, particularly wheat and barley. Fusarium Head Blight causes considerable annual losses worldwide, with severe outbreaks reported in several countries such as the United States (McMullen et al. 2012), Canada (Xue et al. 2019), and across Europe (Figueroa et al. 2018; Parry et al. 1995). Beyond crop damage, *F. graminearum* poses a serious health threat to humans and animals due to its production of mycotoxins, inducing diverse toxic effects on the liver, kidney, immune, and nervous system (Escrivá et al. 2015; Ferrigo et al. 2016; Placinta et al. 1999). Stringent regulatory thresholds on mycotoxin levels have thus been set up in cereal grains and cereal-based products (i.e., EU regulation 2023/915). The most important mycotoxins produced by *F. graminearum* in grains include deoxynivalenol (DON) and its acetylated derivatives (15-acetyldeoxynivalenol, 15ADON; and 3-acetyldeoxynivalenol, 3ADON), nivalenol (NIV), and zearalenone (ZEA).

To develop adequate control strategies for preventing FHB epidemics, it is critical to track *Fusarium* inoculum sources at both species and population levels (Sneideris et al. 2020). Among the various *Fusarium* species identified as causal agents of FHB, *F. graminearum* was reported as the most predominant in wheat grains and maize residues in France (Cobo-Díaz et al. 2019; Nguyen et al. 2024b). *Fusarium graminearum* can survive saprophytically on maize residues and colonize the wheat heads primarily during flowering (Leplat et al. 2013). The implementation of conservation tillage practices, such as no to minimum tillage, has substantially increased in different countries over recent decades to help prevent soil erosion and degradation (Holland 2004; Kassam et al. 2019). Unlike tillage, these practices result in the presence of crop residues on the soil surface, which are considered the primary inoculum sources for FHB causal agents (Leplat et al. 2013). Of all preceding crops, maize presents the highest risk because of the large amount of maize residues after harvest and its chemical composition (low C/N), which provide a favorable habitat allowing *F. graminearum* winter survival (Leplat et al. 2013). Although *F. graminearum* is homothallic and capable of self-fertilization, evidence suggests that in nature, *F. graminearum* populations outcross frequently (Fulcher et al. 2019; Gardiner et al. 2020). The ascospores and conidia produced during sexual and asexual phases on crop residues are considered the primary inoculum sources for wheat head infection (Osborne and Stein 2007). Under conducive conditions during spring, spores produced by *Fusarium* species are expelled and dispersed over long distances by air movements and/or spread through rain (Fernando et al. 2000; Paul et al. 2004).

Population genetics and genomics contribute to understanding the epidemiology of plant diseases by enabling the tracking of inoculum sources and dispersal trajectory, as well as the evaluation of potential for host jumps and mating (Nguyen et al. 2024a). For instance, high genetic variation in a population associated with linkage disequilibrium and the presence of opposite mating-type genes for heterothallic fungi serve as an indicator of frequent sexual reproduction and the potential for rapid evolution. Genetic population analysis also provides insights into gene flow and migration of genes and genotypes between populations (Milgroom and Fry 1997). In *F. graminearum* populations, most studies revealed high genetic diversity even at a very small field scale, which is linked to high gene flow between populations and frequent recombination events within the population (reviewed in Nguyen et al. 2024a). However, *Fusarium* population genetic studies have mainly focused on studying populations isolated from grains at various spatio-temporal scales. To the best of our knowledge, a comparison of the *Fusarium* populations originating from the primary inoculum sources, that is crop residues, has yet to be undertaken. Such a study may help determine the extent to which residues contribute to wheat head contamination.

In this context, the objective of the present work was to evaluate the chemotype distribution, clonal composition, and genetic variation across *F. graminearum* populations originating from different geographical locations and sources. Additionally, the spatial dispersion of inoculum from maize residues to wheat grains was investigated to better understand the role of maize residues in wheat head infection. To do so, a collection of 588 single-spore *F. graminearum* isolates was built. Isolates were collected in four wheat fields over two growing seasons (2020 to 2022) and originated from two substrates: wheat grains (primary host - 146 isolates) and maize residues (potential primary inoculum source - 442 isolates). A sequence-based microsatellite genotyping method (SSRseq) using 34 SSR markers was used to assess the genetic diversity and structure of *F. graminearum* populations, while the chemotype distribution was evaluated by multiplex PCR analyses.

## Materials and Methods

### Sampling of *Fusarium graminearum* populations

Four wheat fields (referred to as F1, F3, F4, and F6 with F4 and F6 located in the same city) located in Brittany, France, were selected for sampling *F. graminearum* populations across two consecutive wheat cycles: 2020–2021 (Y1) and 2021–2022 (Y2) (Figure 1). All wheat fields were cultivated under a maize-wheat rotation system with minimum tillage, except for F1Y1, which was ploughed under exceptional circumstances. Geographical distances from F1 to F3, F1 to F4&F6, and F3 to F4&F6 are 90, 122, and 70 km, respectively. In Y1, moderate (F1 and F4) to severe (F3 and F6) FHB infections were observed in these fields, resulting in the mycotoxin contamination of harvested grains, particularly with DON. In contrast, low to no FHB infection was detected on wheat grains in Y2 (Nguyen et al. 2024b).

**Figure 1.**
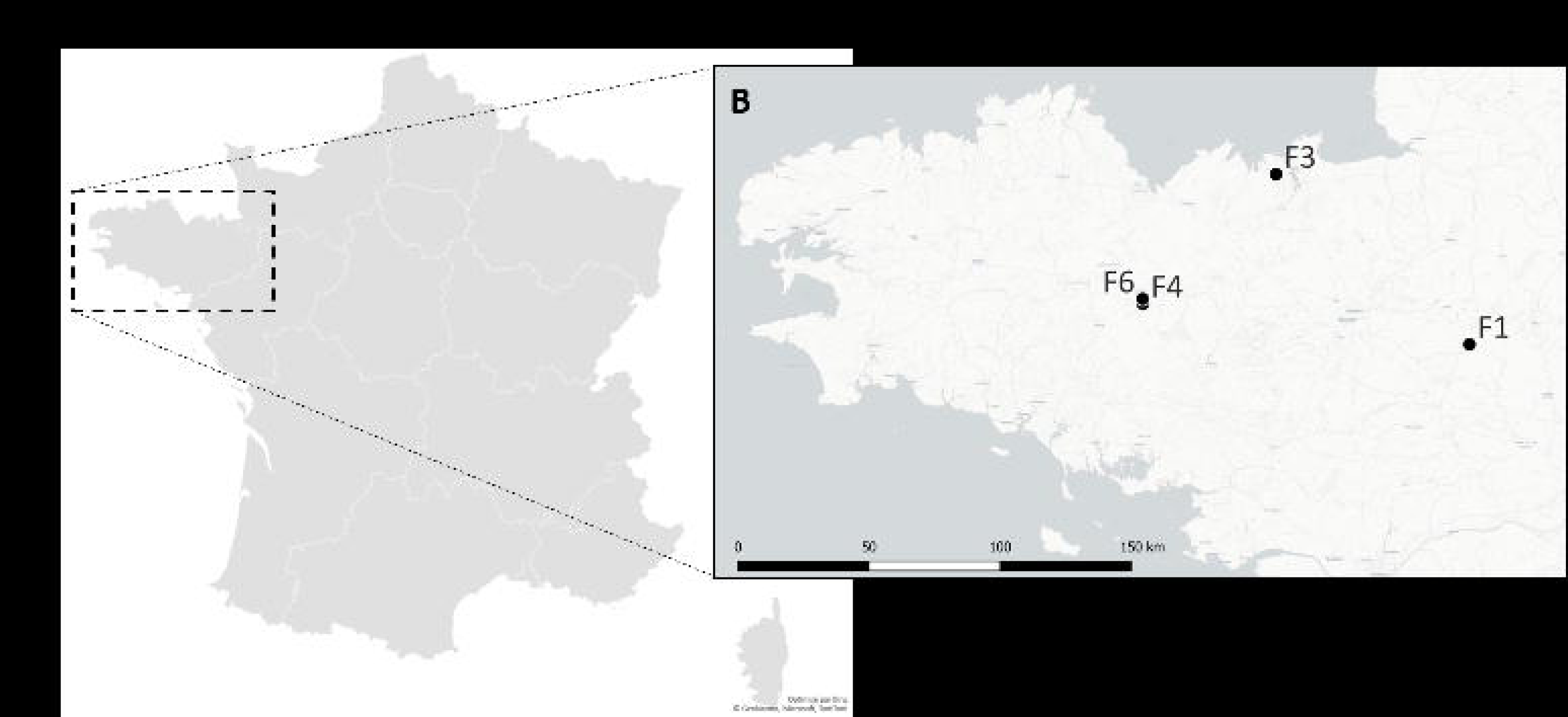
Geographic locations of four wheat fields sampled during the 2020-2021 wheat cycle. (A) French map. (B) Zoomed-in view of the Brittany region in northwest France. Black dots represent the sampled wheat fields.

Sampling was conducted on maize residues and wheat grains to assess *F. graminearum* populations. Maize residues were collected in both Y1 and Y2 at four time points: after the maize harvest (T1), at the seedling stage (T2), 2–3 weeks after wheat flowering (T3), and at harvest (T4), except for F1, where only the T1 sampling in Y1 was conducted. Wheat grains were sampled exclusively in Y1 at two time points: 2–3 weeks after flowering (T3) and at wheat harvest (T4), as low to no FHB infection was observed in Y2. In total, twelve field × substrate × year populations were defined based on the combination of field, substrate, and year.

Three sampling points per field were selected, with each sampling point consisting of a composite of 10 residue subsamples and 30 wheat heads collected within a 5-m radius half-circle. Samples were stored at 4°C before culturing on potato dextrose agar (PDA; BD Difco, Sparks, MA, USA), with maize residues stored in zip-lock bags for up to 3 weeks and wheat heads in paper bags for up to one month. Due to time constraints, maize residues from T3Y1 and T4Y1 were stored in glycerol 20% at −80°C before *F. graminearum* isolation.

Each maize residue subsample (corresponding to one sampling point per field) was previously cut into pieces of 0.5 cm, surface-disinfected by immersion in a 1% (v/v) sodium hypochlorite solution for 1 minute, followed by two rinses with sterile demineralized water, while keeping subsamples separated from each other. The surface-disinfected samples were aseptically dried in sterile filter papers for at least 15 min, then five pieces of each residue were plated on PDA medium, supplemented with a mixture of 50 mg L^−1^ of streptomycin, penicillin, and chlortetracycline (Sigma-Aldrich, MO, USA) (hereafter referred to as PDA+ATB). For wheat grain samples collected at T3, five grains from each head were directly plated on PDA+ATB medium for *F. graminearum* isolation. At T4, both surface-disinfected and non-disinfected grains were used with the same processes as described above. The maize residues and wheat grains on PDA+ATB medium were incubated at room temperature for 5–7 days. One isolate from each plate, morphologically identified as *F. graminearum*, was purified and stored on PDA at 4°C.

For single-spore isolation, collected isolates were grown for 5–7 days at room temperature. Conidia were washed from the cultures using 5 mL of sterile demineralized water supplemented with Tween 80 (2 drops in 1 L). The spore suspension was streaked on PDA and incubated overnight at room temperature. A single germinating conidium was transferred onto PDA and subsequently stored at 4°C.

### DNA extraction

To extract genomic DNA, single spore isolates were cultivated on PDA plates and grown for 7 days at room temperature. Fungal tissues were scraped with a sterile scalpel and placed into tubes containing type A beads (Macherey-Nagel MN, Düren, Germany) and stored at −20°C until DNA extraction. Genomic DNA was automatically extracted using KingFisherTM Duo Prime (Thermo Fisher Scientific, MA, USA) and Mag-Bind^®^ Universal Pathogen DNA 96 Kit (Omega Bio-Tek, GA, USA).

### Molecular identification of *Fusarium graminearum*

The identity of isolates was subsequently verified by PCR using two species-specific primer pairs: Fg16NF (5’-ACAGATGACAAGATTCAGGCACA-3’) and Fg16NR (5’-TTCTTTGACATCTGTTCAACCCA-3’) (Nicholson et al. 1998), then FgTEFf124 (5’-CGGTCACTTGATCTACCAG-3’), FgTEFr590 (5’-GAATGTGATGACAGCAGTG-3’), and Fg7TEFf364 (5’-CTCGAGCGACAGGCGTC-3’) (Yang et al. 2008). PCR was performed using the GoTaq^®^ kit and dNTP mix from Promega, WI, USA, under the conditions reported in the cited studies with a few modifications (Table 1). Amplicons were analyzed by electrophoresis through 1% agarose gels with Midori Green Advance^®^ stain (Nippon Genetics Co. Europe GmbH, Düren, Germany) in 1X TAE buffer (Tris-acetate-EDTA, Promega), then visualized under UV light. Isolates, for which DNA was not amplified in one of the two PCR runs, were subsequently verified by analysis of translation elongation factor 1L (*EF1*L) sequences using primer pairs EF1f and EF1r (Brygoo and Gautier 2007). Sequencing was performed by Eurofins Genomics (Ebersberg, Germany). The obtained sequences were analyzed using the Geneious software (Kearse et al. 2012) and subjected to a BLASTn search of the NCBI database.

**Table 1.**
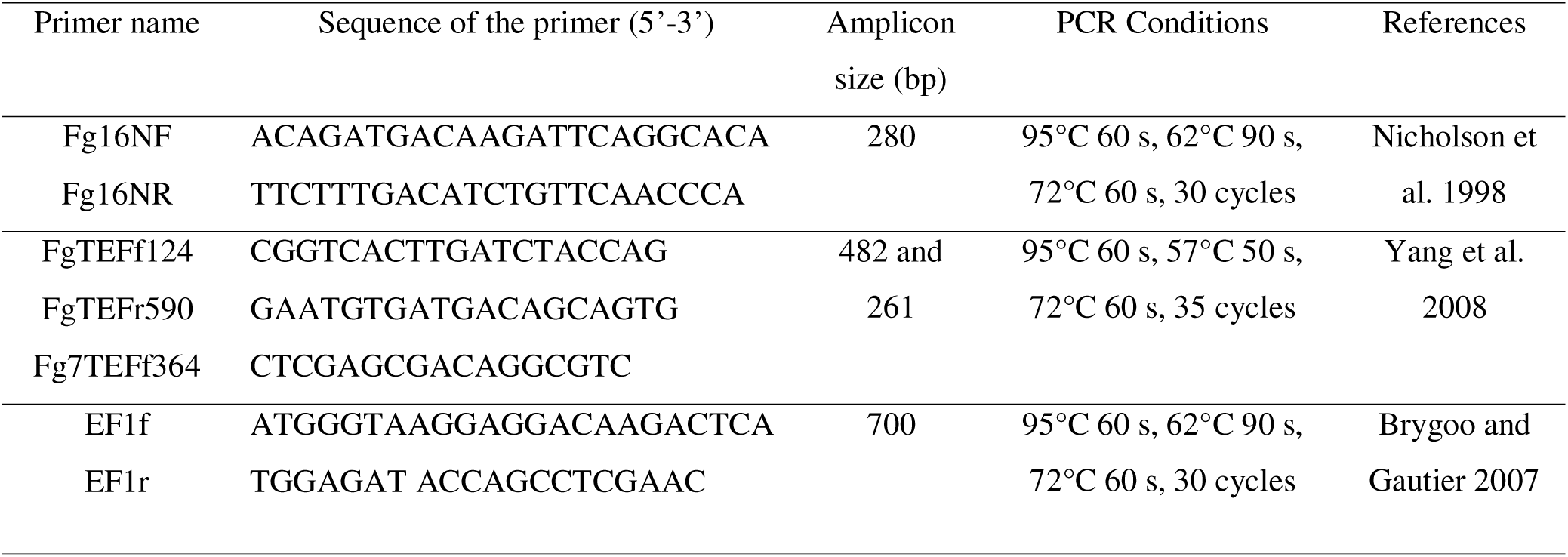
Primer sequences, annealing temperatures, and sizes of amplicon fragments used to taxonomically assign/confirm *Fusarium graminearum* isolates from maize residues and wheat grains.

### Sequence-based microsatellite genotyping

Simple sequence repeat (SSR) genotyping, including whole-genome SSR detection, primer design, SSR sequencing, and bioinformatic analyses, was performed at the Genome Transcriptome Facility of Bordeaux (PGTB, Bordeaux, France). The workflow details are provided in Lepais et al. (2020). Briefly, the QDD version 3.1.2 pipeline (Meglécz et al. 2014) was employed for SSR detection and primer design using the *F. graminearum* reference genome sequence from NCBI (accession numbers GCA_020991245.1). Sixty out of 6626 candidate microsatellites were selected based on rigorous quality criteria as described by Lepais et al. (2020), and subsequently tested for amplification in simplex PCR, followed by amplification of 55 reliable loci in multiplex PCR. Amplicons were indexed and sequenced. Two separate SSR genotyping runs were performed. The first run included *F. graminearum* isolates collected from residues (T1Y1, T2Y1) and grains (T3Y1, T4Y1), sequenced using Illumina NextSeq 2000 in a paired-end 2 x 250 bp configuration. The second run included isolates from residues (T3Y1, T4Y1, T1Y2 to T4Y2), sequenced using iSeq100 in a paired-end 2 x 150 bp configuration. Sequences from two runs were combined for a common analysis thanks to anchored samples. Data analysis was conducted using the bioinformatic pipeline outlined by Lepais et al. (2020), which integrated the FDSTools version 1.2.0 pipeline (Hoogenboom et al. 2017). This pipeline allowed the generation of complete genotypic profiles for each isolate, encompassing allelic polymorphism across all tested 55 loci, as recommended by Lepais et al. (2020). Additionally, sequences from 95 *F. graminearum* isolates of the first run were amplified and sequenced in duplicate to check the reproducibility as well as to determine the allelic error rate and overall missing data rate for each locus.

### SSR quality filtering

Loci showing more than 20% missing data and/or 3% allelic error were excluded from the analysis (3 loci). Additionally, individuals with more than 20% heterozygosity were discarded, considering the haploid nature of our isolates (3 isolates). Genotypes of repeated individuals were compared, and if discrepancies were detected between repeated isolates, the genotype was assigned as missing data (NA). One representative genotype was kept for each repeated isolate to ensure data consistency. Following this verification step, monomorphic loci (1 locus) and those exceeding 20% of missing data were discarded (that is, 0 loci), as well as isolates with over 20% of missing data (7 isolates). The polymorphism information content (PIC) was calculated for each SSR marker using the poppr R package version 2.9.4, and markers with PIC values below 0.25 were discarded due to low informativeness (17 loci) (Serrote et al. 2020). Overall, among the 55 SSRs and 598 individuals initially tested, 34 polymorphic markers and 588 individuals were selected for further genetic diversity analysis.

### Population genetic analysis

#### Analysis of genetic diversity

GenAlEx version 6.5 software (Peakall and Smouse 2012) was used to calculate several parameters of genetic diversity, including the number of private alleles, mean number of different alleles (Na), effective number of alleles (Ne, Kimura and Crow 1964), Shannon’s information index of allelic diversity (I, Brown and Weir 1983), and Nei’s unbiased gene diversity (uh, Nei 1978). In addition, genotype richness and diversity within populations were estimated using the poppr R package version 2.9.4 (Kamvar et al. 2014). The genotypic diversity indices included the number of multilocus genotypes (MLGs), number of expected MLGs at the smallest sample size (eMLGs), Shannon-Wiener index of MLG diversity (H, Shannon and Weaver 1964), Simpson’s index (λ, Simpson 1949), evenness of MLGs (E.5, distribution of genotypic abundance, Grünwald et al. 2003). The E.5 value, ranging from 0 (uneven) to 1 (equal distribution of genotypes), measures the distribution of genotypic abundances (Grünwald et al. 2003). Pairwise comparisons of populations were analyzed using a t-test of the online GRAPHPAD QUICKCALCS software (https://www.graphpad.com/quickcalcs/ttest1.cfm?Format=SD) to compare the values of these parameters. To measure the degree of association among alleles, multilocus linkage disequilibrium (LD) was estimated on SSR data with the poppr R package (Kamvar et al. 2014), using both non-clone-corrected and clone-corrected data (where a single representative individual of each MLG is kept for analysis). Recombination evidence was assessed based on the indices of association I_A_ (Brown et al. 1980) and rbarD (Agapow and Burt 2001) with 1000 permutations.

#### Analysis of genetic distribution and differentiation

To visualize the genetic relationship among genotypes, a principal coordinate analysis (PCoA) based on Nei’s unbiased genetic distance (Nei 1978) was conducted. Genetic variation among and within populations was estimated using analysis of molecular variance (AMOVA), with significance determined through 999 permutations. Both PCoA and AMOVA analyses were performed using GenAlEx software (Peakall and Smouse 2012). From AMOVA, the genetic differentiation PhiPT (or Φ_PT_, equivalent to F_ST_ fixation index for haploid organisms) among all pairwise populations and gene flow estimation (Nm, indicating the number of migrants) were obtained. PhiPT values have been used to calculate gene flow as Nm = (1/PhiPT - 1)/4.

Population structure of the clone-corrected dataset was inferred using the model-based clustering program Structure version 2.3.4 (Pritchard et al. 2000). To determine the optimal number of clusters (K), we tested a series of K values from 1 to 10, performing 15 independent runs for each K with a burn-in of 100,000 and run length of 1,000,000 iterations. The most likely number of clusters was estimated using the method proposed by Evanno et al. (2005), which involves plotting the distribution of ΔK, an ad hoc quantity based on the second-order rate of change of the mean likelihood of K. ΔK was estimated using the online application Structure Harvester (Earl and vonHoldt 2012), then structure results were visualized with the CLUMPAK web-based software (https://tau.evolseq.net/clumpak/) (Kopelman et al. 2015). The discriminant analysis of principal components (DAPC) was applied to explore genetic relationships among pre-defined groups of isolates and patterns of population structure (Jombart et al. 2010) using the adegenet R package version 2.1.10.

The global optimal eBURST algorithm (goeBURST), implemented in Phyloviz software version 2.0 (Francisco et al. 2009), was used to assign clusters of related MLGs (hereafter referred to as CCs for clonal complexes), at the triple locus variant (TLV) level of clone-corrected data. Clonal complexes were defined as groups of MLGs sharing at least 31 out of 34 identical alleles with at least one other MLG. A central MLG involved in the highest number of relationships with TLVs was identified and allowed to determine the CC, including this central MLG, its neighbors, and the neighbors’ neighbors. To infer the between-population relationships, a minimum spanning genotype network was constructed using the poppr R package based on relative dissimilarity distances of non-clone-corrected data.

### Identification of trichothecene and zearalenone chemotypes

Trichothecene and zearalenone chemotypes of each *F. graminearum* strain were identified using multiplex PCR, as described by Ward et al. (2008) and Sim et al. (2018), respectively. For trichothecene chemotyping, TRI3 multiplex PCR with primers 3CON, 3NA, 3D3A, and 3D15A was employed, generating an 840 bp fragment for the NIV chemotype, 610 bp for the 15ADON chemotype, and 243 bp for the 3ADON chemotype (Ward et al. 2008). For strains that did not yield results with TRI3, TRI12 multiplex amplification was performed using primers 12CON, 12NF, 12-15F, 12-3F, generating fragments of 840bp (NIV), 670 bp (15ADON), and 410 bp (3ADON) (Ward et al. 2008). The thermal cycling conditions for TRI3 amplification were: 94°C for 4 min, followed by 35 cycles of 94°C for 1 min, 58°C for 40 s, 72°C for 40 s, and a final extension at 72°C for 6 min (Oghenekaro et al. 2021). The thermal cycling conditions for TRI12 amplification were: 94°C for 2 min, followed by 30 cycles of 94°C for 30 s, 52°C for 30 s, 72°C for 1 min, and a final extension at 72°C for 10 min (Ward et al. 2008). PCR amplicons were separated on 1.5% (m/v) agarose gel in 1X TAE buffer (Tris-acetate EDTA, Promega), stained with Midori Green Advance^®^ (Nippon Genetics Co. Europe GmbH, Düren, Germany), and sizes were estimated with a 100 bp DNA ladder (Promega).

For the identification of ZEA-producing strains, primer pairs targeting the *PKS3* (280 bp), *PKS13* (192 bp), *ZEB1* (129 bp), and *ZEB2* (80 bp) genes were used (Sim et al. 2018). The thermal cycling conditions used were: 94°C for 4 min, followed by 35 cycles of 94°C for 1 min, 65°C for 30 s, 72°C for 1 min, and a final extension at 72°C for 10 min. PCR amplicons were separated on 3% (m/v) agarose gel in 1X TAE buffer (Promega), stained with Midori Green Advance^®^, and sizes were estimated with a 100 bp DNA ladder (Promega).

## Results

### Sample collections

A total of 858 isolates were initially collected based on macro- and microscopic morphological characteristics. Molecular-based taxonomic assignment to *F. graminearum* was confirmed for 598 isolates, with the misidentifications mainly being due to *F. cerealis* (synonymous with *F. crookwellense*), morphologically similar to *F. graminearum* (e.g., similar shape of macroconidia, absence of microconidia, red pigmentation) (Amarasinghe et al. 2015). Finally, a total of 450 *F. graminearum* isolates from distinct maize residues (215 from Y1 and 235 from Y2) and 148 isolates from wheat heads (from Y1) were used for genetic structure analysis and assessment of chemotype diversity. After filtering SSR genotyping data for quality, 588 isolates were retained for further analysis.

### Population genetic diversity and structure according to geographic locations

In total, 453 MLGs were obtained after genotyping 588 isolates using 34 SSR markers (Table 2). Of the 453 MLGs, 395 were represented by a single isolate (Figure S1). The most frequent MLGs were shared by 21 (MLG 167) and 7 isolates (MLG 244), all of which were collected from both maize residues and wheat grains (Figure 2). The most frequent MLG was found across nine out of the twelve field × substrate × year populations. The number of eMLG (number of MLGs normalized by sample size) was similar between the twelve populations (*p* > 0.05). In addition, the genotype distribution (E.5) showed that most populations had an even distribution of MLGs, except F3-grains-Y1. The Shannon-Wiener (H) and Simpson’s index (λ) were relatively similar across the twelve populations, ranging from 3.108 to 4.054 and from 0.941 to 0.982, respectively.

**Figure 2.**
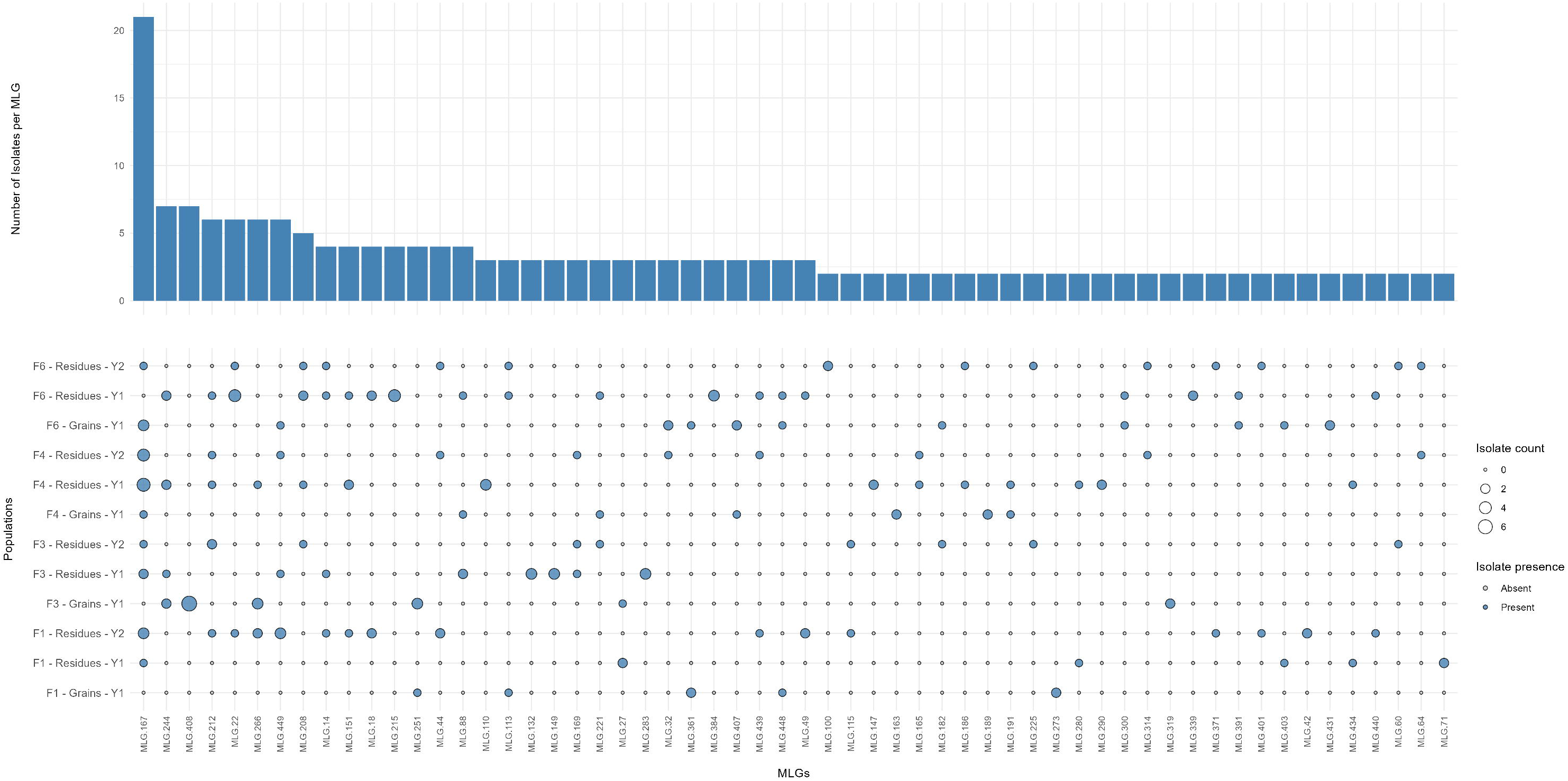
Multilocus genotypes (MLGs) of twelve *Fusarium graminearum* populations from four different wheat fields and two types of substrates (wheat grains and maize residues) across two wheat cycles in Brittany, France, based on sequence-based microsatellite genotyping of 34 markers. The barplot represents the number of isolates per MLG. The bubble plot depicts the presence or absence of isolates across 12 populations. Unique MLGs (represented by one individual) were discarded from this plot (395 MLGs).

**Table 2.** Genetic diversity of twelve field × substrate × year populations of *Fusarium graminearum* in Brittany, France, based on sequence-based microsatellite genotyping of 34 markers.

| Substrate | Field | Year | Pop | N | MLG | eMLG | SE | H | □ | E.5 |
| --- | --- | --- | --- | --- | --- | --- | --- | --- | --- | --- |
| All populations |  |  |  | 588 | 453 | 24.331 | 0.844 | 5.919 | 0.996 | 0.635 |
| Grains | F1 | Y1 | F1 - Grains - Y1 | 26 | 24 | 23.154 | 0.361 | 3.151 | 0.956 | 0.963 |
|  | F3 | Y1 | F3 - Grains - Y1 | 41 | 29 | 19.236 | 1.438 | 3.153 | 0.941 | 0.713 |
|  | F4 | Y1 | F4 - Grains - Y1 | 32 | 30 | 23.79 | 0.671 | 3.379 | 0.965 | 0.968 |
|  | F6 | Y1 | F6 - Grains - Y1 | 47 | 42 | 23.477 | 0.99 | 3.692 | 0.973 | 0.932 |
| Residues |  | Y1 | F1 - Residues - Y1 | 25 | 23 | 23 | 0 | 3.108 | 0.954 | 0.961 |
|  | F1 | Y2 | F1 - Residues - Y2 | 59 | 50 | 23.213 | 1.126 | 3.848 | 0.977 | 0.914 |
|  |  | Y1 | F3 - Residues - Y1 | 55 | 47 | 23.041 | 1.167 | 3.777 | 0.975 | 0.897 |
|  | F3 | Y2 | F3 - Residues - Y2 | 54 | 53 | 24.79 | 0.407 | 3.963 | 0.981 | 0.989 |
|  |  | Y1 | F4 - Residues - Y1 | 67 | 57 | 23.144 | 1.201 | 3.953 | 0.978 | 0.851 |
|  | F4 | Y2 | F4 - Residues - Y2 | 58 | 55 | 24.179 | 0.827 | 3.965 | 0.979 | 0.910 |
|  |  | Y1 | F6 - Residues - Y1 | 65 | 53 | 22.696 | 1.297 | 3.868 | 0.976 | 0.854 |
|  | F6 | Y2 | F6 - Residues - Y2 | 59 | 58 | 24.825 | 0.38 | 4.054 | 0.982 | 0.990 |

N = number of isolates, MLG = number of observed multilocus genotype, eMLG = number of expected MLGs at the smallest sample size based on rarefaction, SE = standard error based on
eMLG, H = Shannon-Wiener index, $\lambda$ = Simpson's index, E.5 = evenness (distribution of genotype abundance in a sample evenness).

The number of different alleles for each locus was more than three within the twelve populations (Table 3). Private alleles (number of alleles unique to a single population) were detected in all populations, ranging from 0.029 to 0.265, except for F4 grains, where no private allele was found. The average values of Shannon’s information index and observed haploid diversity ranged from 0.281 to 0.428 and from 0.162 to 0.227, respectively. No significant difference in allelic diversity was found across the twelve field × substrate × year populations.

**Table 3.**
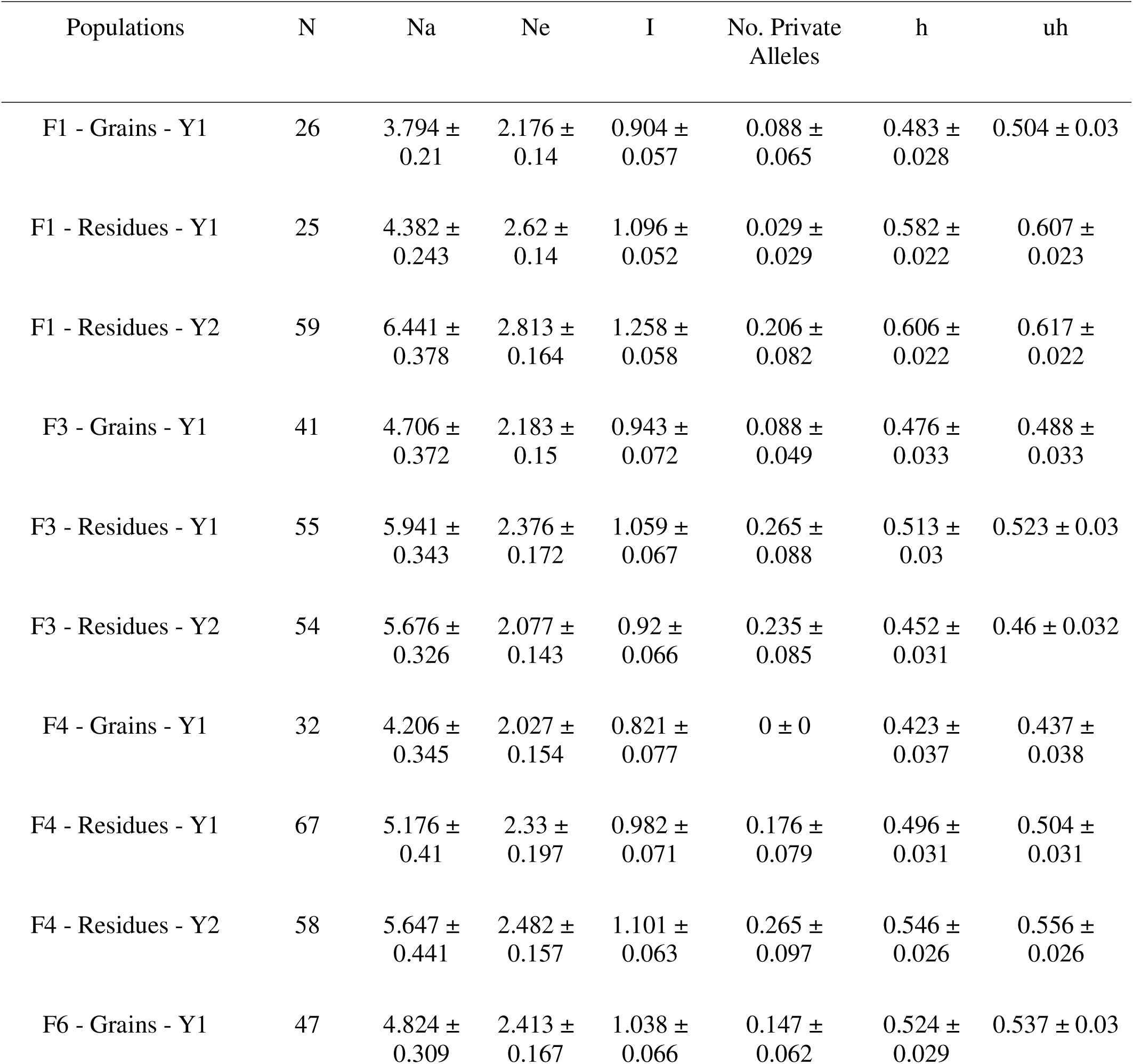
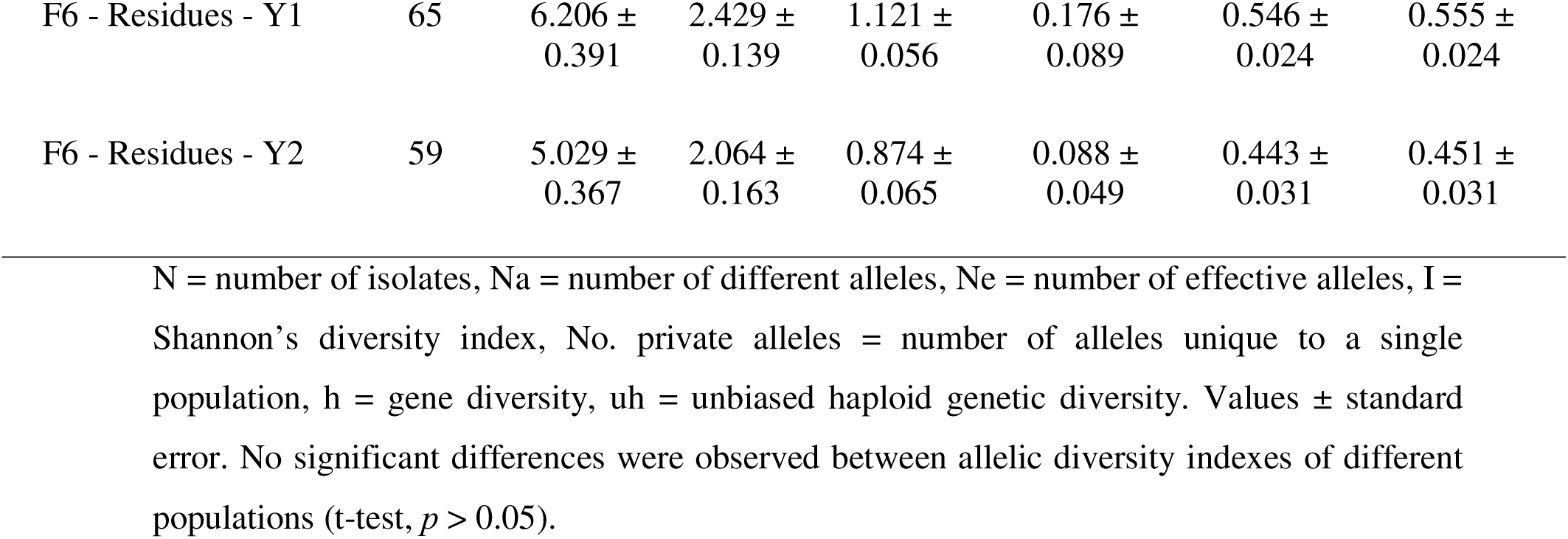
Allelic diversity within the twelve field × substrate × year populations of *Fusarium graminearum* in Brittany, France, based on sequence-based microsatellite genotyping of 34 markers.

| Populations | N | Na | Ne | I | No. Private Alleles | h | uh |
| --- | --- | --- | --- | --- | --- | --- | --- |
| F1 - Grains - Y1 | 26 | 3.794 ± 0.21 | 2.176 ± 0.14 | 0.904 ± 0.057 | 0.088 ± 0.065 | 0.483 ± 0.028 | 0.504 ± 0.03 |
| F1 - Residues - Y1 | 25 | 4.382 ± 0.243 | 2.62 ± 0.14 | 1.096 ± 0.052 | 0.029 ± 0.029 | 0.582 ± 0.022 | 0.607 ± 0.023 |
| F1 - Residues - Y2 | 59 | 6.441 ± 0.378 | 2.813 ± 0.164 | 1.258 ± 0.058 | 0.206 ± 0.082 | 0.606 ± 0.022 | 0.617 ± 0.022 |
| F3 - Grains - Y1 | 41 | 4.706 ± 0.372 | 2.183 ± 0.15 | 0.943 ± 0.072 | 0.088 ± 0.049 | 0.476 ± 0.033 | 0.488 ± 0.033 |
| F3 - Residues - Y1 | 55 | 5.941 ± 0.343 | 2.376 ± 0.172 | 1.059 ± 0.067 | 0.265 ± 0.088 | 0.513 ± 0.03 | 0.523 ± 0.03 |
| F3 - Residues - Y2 | 54 | 5.676 ± 0.326 | 2.077 ± 0.143 | 0.92 ± 0.066 | 0.235 ± 0.085 | 0.452 ± 0.031 | 0.46 ± 0.032 |
| F4 - Grains - Y1 | 32 | 4.206 ± 0.345 | 2.027 ± 0.154 | 0.821 ± 0.077 | 0 ± 0 | 0.423 ± 0.037 | 0.437 ± 0.038 |
| F4 - Residues - Y1 | 67 | 5.176 ± 0.41 | 2.33 ± 0.197 | 0.982 ± 0.071 | 0.176 ± 0.079 | 0.496 ± 0.031 | 0.504 ± 0.031 |
| F4 - Residues - Y2 | 58 | 5.647 ± 0.441 | 2.482 ± 0.157 | 1.101 ± 0.063 | 0.265 ± 0.097 | 0.546 ± 0.026 | 0.556 ± 0.026 |
| F6 - Grains - Y1 | 47 | 4.824 ± 0.309 | 2.413 ± 0.167 | 1.038 ± 0.066 | 0.147 ± 0.062 | 0.524 ± 0.029 | 0.537 ± 0.03 |
| F6 - Residues - Y1 | 65 | 6.206 ±<br>0.391 | 2.429 ±<br>0.139 | 1.121 ±<br>0.056 | 0.176 ±<br>0.089 | 0.546 ±<br>0.024 | 0.555 ±<br>0.024 |
| F6 - Residues - Y2 | 59 | 5.029 ±<br>0.367 | 2.064 ±<br>0.163 | 0.874 ±<br>0.065 | 0.088 ±<br>0.049 | 0.443 ±<br>0.031 | 0.451 ±<br>0.031 |

N = number of isolates, Na = number of different alleles, Ne = number of effective alleles, I = Shannon's diversity index, No. private alleles = number of alleles unique to a single population, h = gene diversity, uh = unbiased haploid genetic diversity. Values ± standard error. No significant differences were observed between allelic diversity indexes of different populations (t-test, $p > 0.05$ ).

An AMOVA was performed to explore the impact of geographic origin on genetic structure and showed that the majority of genotypic diversity (99%) was found within each geographical area rather than among (1%) (Table 4). In agreement with this result, differentiation of populations by geographic location revealed significant but small pairwise differences between populations (PhiPT < 0.1). Likewise, DAPC analysis revealed no significant differentiation based on the geographic origin of isolates (Figure 3A).

**Figure 3.**
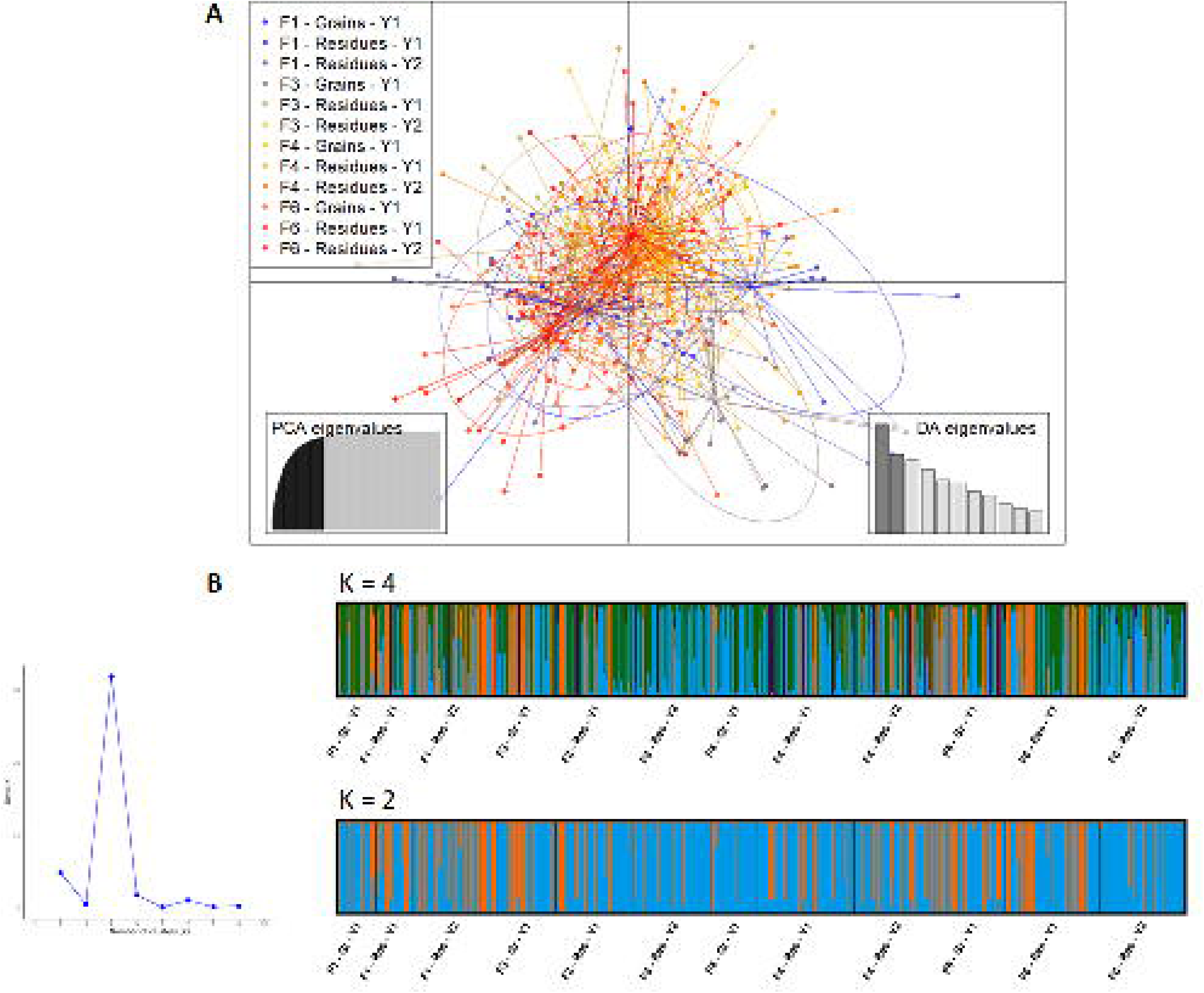
Population structure analysis of *Fusarium graminearum* populations based on fields and substrates from four wheat fields in Brittany, France. (A) Discriminant analysis of principal components (DAPC) for 588 *F. graminearum* isolates using a dataset of 34 microsatellite markers. The axes represent the first two linear discriminants. (B) Inferred population genetic structure of *F. graminearum* populations (K = 2 and K = 4) based on 34 microsatellite markers using STRUCTURE software 2.3.4.

**Table 4.** Analysis of molecular variance (AMOVA) of *Fusarium graminearum* populations from three geographic areas in Brittany, France.

| Source | df | SS | MS | Est. Var. | % | Overall<br>PhiPT | Probability |
| --- | --- | --- | --- | --- | --- | --- | --- |
| Among<br>populations | 2 | 47665.093 | 23832.546 | 79.929 | 1% | 0.008 | 0.001 |
| Within populations | 585 | 5848402.276 | 9997.269 | 9997.269 | 99% |  |  |
| Total | 587 | 5896067.369 |  | 10077.198 | 100% |  |  |
| Pairwise population PhiPT values |  |  |  |  |  |  |  |
|  | F1 | F3 | F4&F6 |  |  |  |  |
| F1 | 0.000 | 0.001 | 0.007 |  |  |  |  |
| F3 | 0.016 | 0.000 | 0.097 |  |  |  |  |
| F4&F6 | 0.010 | 0.003 | 0.000 |  |  |  |  |

df = degree of freedom, SS = sum of squares, MS = mean squares, Est. Var. = estimate of variance, % = percentage of total variation, Overall PhiPT = genetic differentiation among populations. Probability based on 999 permutations. Pairwise population PhiPT values below the diagonal. Probability based on 999 permutations is above the diagonal. Isolates from four wheat fields (F1, F3, F4, and F6) were divided into three geographical areas in Brittany (France), including Gennes-sur-Seiche (F1), Pleslin-Trigavou (F3), and Loudéac (F4&F6).

A Bayesian clustering method, namely STRUCTURE, was used on the clone-corrected dataset to determine population membership of individuals to genetic clusters without prior knowledge of substrate, geographic, or year origins. Only two genetic clusters were identified, both of which were evenly distributed within each geographic population (Figure 3B). The I_A_ and rbarD were used to investigate whether *F. graminearum* populations reproduce clonally (where significant disequilibrium is expected due to linked loci) or sexually (where linkage among loci is not expected). The rbarD value is expected to be 0 where populations are freely recombining (sexual reproduction) and greater than zero if there is an association between alleles (clonality or asexual reproduction). Our results suggested significant linkage disequilibrium (*p* < 0.05) detected in all wheat fields on both non-clone-corrected and clone-corrected datasets (except for F1 using clone-corrected data), indicating that strains were probably under predominant clonal reproduction (Table 5).

**Table 5.** Linkage disequilibrium analysis of *Fusarium graminearum* populations from four wheat fields in Brittany, France.

| Field | I<sub>A</sub> | I<sub>A</sub><sup>a</sup> | r<sub>bar</sub>D | r<sub>bar</sub>D<sup>a</sup> |
| --- | --- | --- | --- | --- |
| F1 | 3.049* | 3.056* | 0.093* | 0.093* |
| F3 | 2.844* | 2.760* | 0.087* | 0.084* |
| F4 | 2.442* | 2.321* | 0.074* | 0.071* |
| F6 | 2.897* | 2.544* | 0.088* | 0.077* |
| All fields | 2.823* | 2.720* | 0.086* | 0.083* |

$I_A$ = index of association, $r_{barD}$ = standardized index of association, \* = significant at $p < 0.001$ , <sup>a</sup> = estimates for $I_A$ and $r_{barD}$ based on clone-corrected dataset. Isolates from each field were collected from maize residues and wheat grains.

### Transfer of *F. graminearum* inoculum from residues to grains

According to AMOVA, the majority of genotypic diversity of *F. graminearum* populations (90%) was found within populations, rather than between populations, based on substrate types (maize residues vs. wheat grains) (Table 6). The proportion of variation among populations was relatively low, accounting for only 10% of the total genetic variance. This low differentiation by substrate type was further illustrated by plotting the genetic distance between isolates as a minimum spanning network, where populations from the two host sources were not clustered separately (Figure 4). Among the 453 MLGs, 15 were found in both substrates, irrespective of geographic locations and sampling years (Figure 2). The minimum spanning network showed high relatedness between genotypes collected in maize residues and wheat grains, with some closely related, similar genotypes present in both substrates (Figure 4).

**Figure 4.**
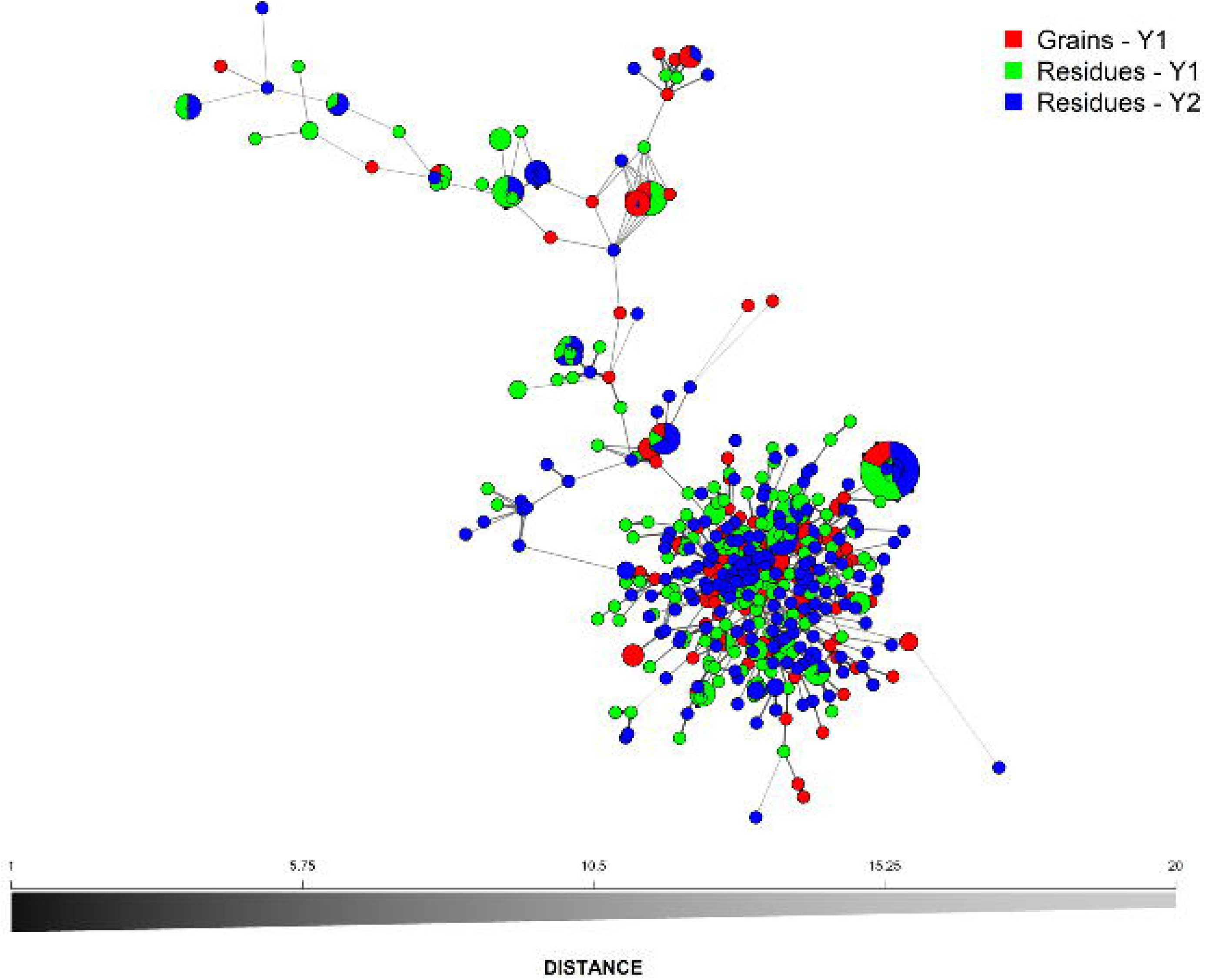
Minimum spanning network based on dissimilarity distances depicting the frequency distribution and the genetic similarity among 453 multilocus genotypes (MLGs) of *Fusarium graminearum* populations according to two types of substrates (maize residues and wheat grains) in Brittany, France. Each node represents a single MLG determined based on 34 microsatellite markers. The node size is the frequency of each genotype sampled. Node colors represent population membership. Edge thickness represents the minimum genetic distance between genotypes.

**Table 6.**
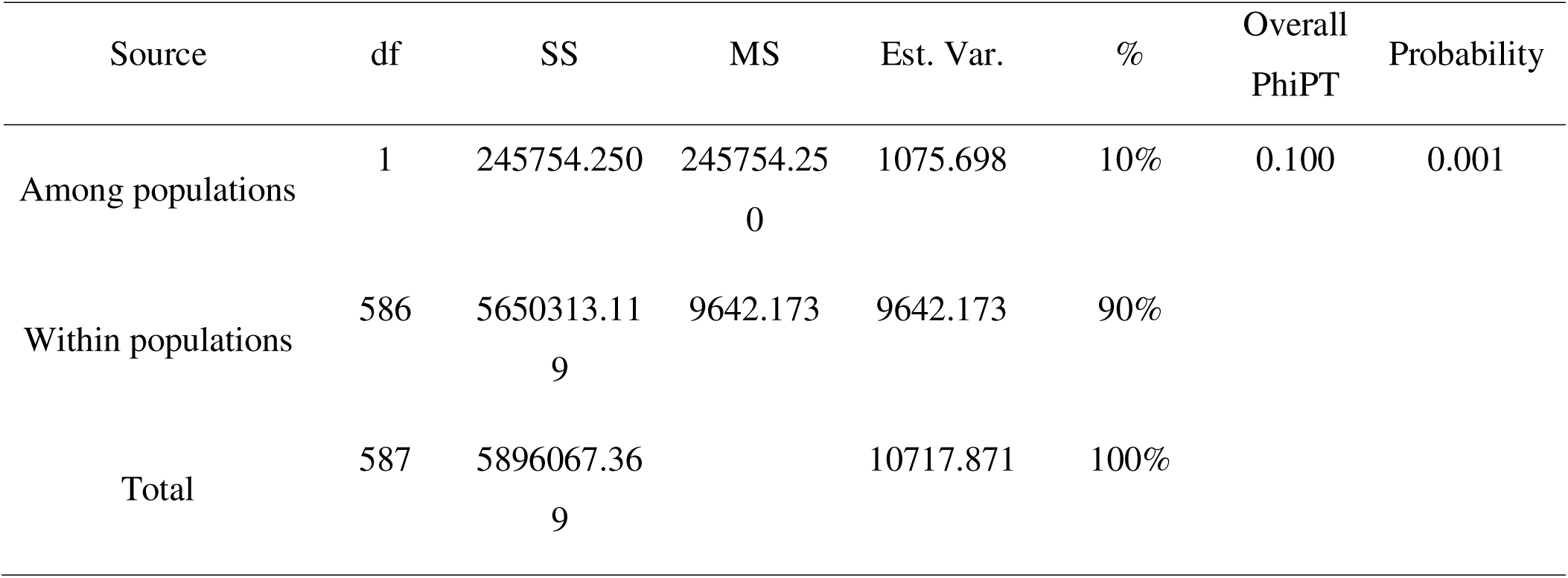
Analysis of molecular variance (AMOVA) of *Fusarium graminearum* populations according to two types of substrates (maize residues and wheat grains) collected in four wheat fields in Brittany, France.

Because of the high genetic diversity of *F. graminearum* populations and the low number of isolates sharing identical SSR profiles across different populations, we clustered them into CCs at the TLV level. The goeBURST analysis identified 335 CCs among 453 MLGs. The most predominant CC was represented by 31 isolates (accounting for 5.3% of isolates) and 8 MLGs (corresponding to 1.8% of MLGs) (Figure S2). The transfer of inoculum from maize residues to wheat grains was estimated by comparing CCs between these two substrates across different fields in Y1 (Figure 5). We found that 3.1 to 29.3% of CCs present in the wheat grain were also detected in residue populations, regardless of the field. Notably, most of the grain-associated CCs in a specific field that were identical to residue-associated CCs of the same field were also found in the residues of all 3 other fields. In other words, there were very few grains and residue-CCs that were uniquely confined to specific fields. Furthermore, a majority of the grain-associated CCs were not found in residues of any fields (F1: 76.9%, F3: 65.9%, F4: 75%, F6: 66.0%).

**Figure 5.**
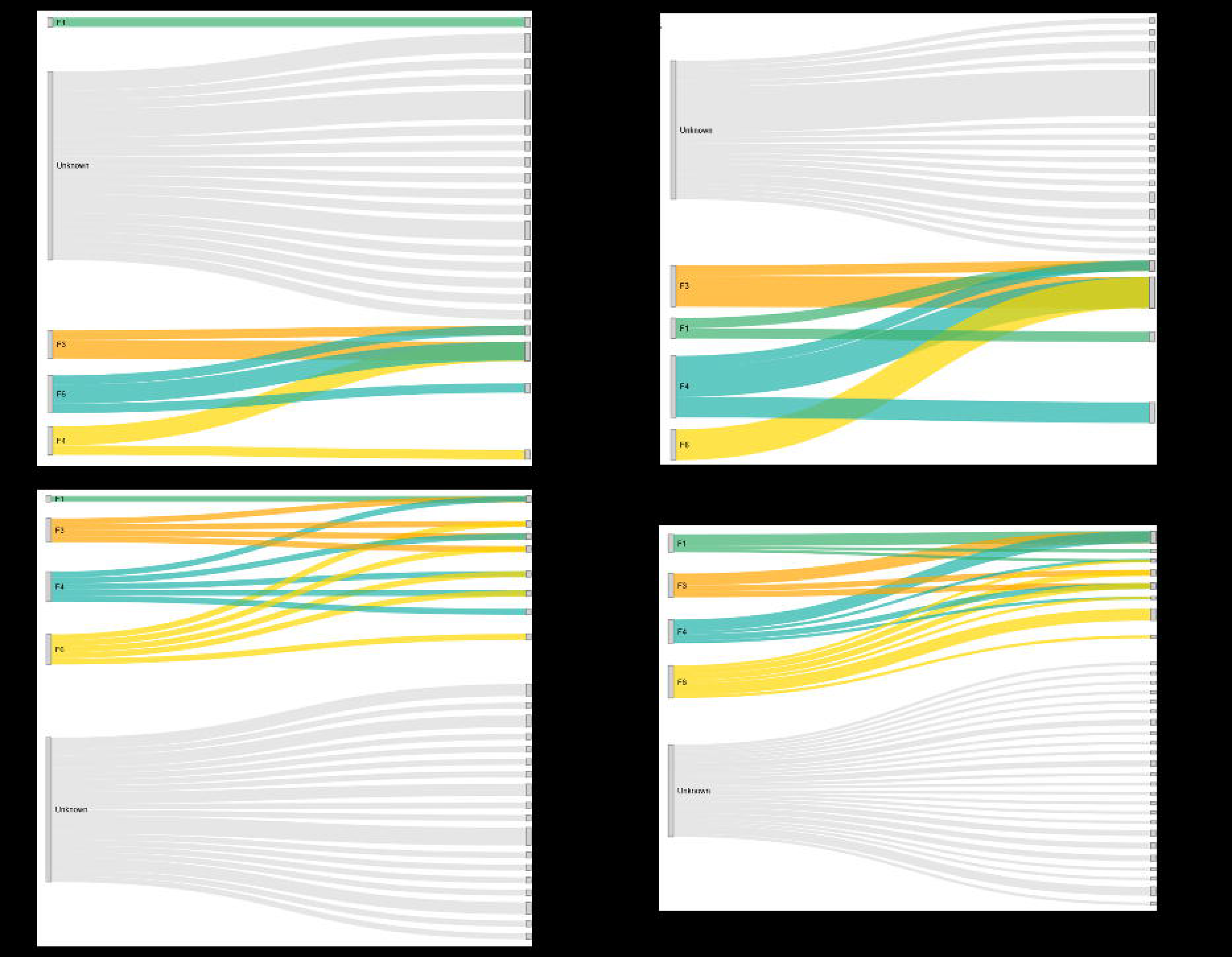
Sankey diagrams illustrating the estimated inoculum transfer of *Fusarium graminearum* populations from maize residues to wheat grains across four wheat fields – F1 (A), F3 (B), F4 (C), and F6 (D) – in Brittany, France. The genetic population dataset was obtained based on sequence-based microsatellite genotyping (SSRseq) of 34 markers. The left side of each diagram represents the potential inoculum sources (maize residues from the four fields and unknown sources). The right side shows the clonal complexes (CCs) of *F. graminearum* populations on wheat grains, identified at single locus variant level using the goeBurst algorithm implemented in the Phyloviz version 2.0 software. The SSR dataset was clone-corrected to include one representative per each MLG for each population (8 populations according to field and substrate). The numbers in brackets indicate the number of MLGs within each CC. The F1 field was tilled with residues only collected at post-harvest of maize before tillage.

### Temporal genetic structure of *Fusarium graminearum* populations in maize residues

To analyze the dynamics of *F. graminearum* populations in residues, isolates collected from four fields and two years were grouped into four temporal groups based on sampling period: early stage (T1T2Y1, T1T2Y2) and later stage of residue degradation (T3T4Y1, T3T4Y2). Each temporal population exhibited high genotypic diversity. According to AMOVA, the majority of genotypic diversity of *F. graminearum* populations (99%) was found within sampling time points, while only a small fraction was attributed to differences between sampling stages (Table 7). The LD values were very low across all temporal groups, with rD varying from 0.073 to 0.093 (*p* < 0.001).

**Table 7.**
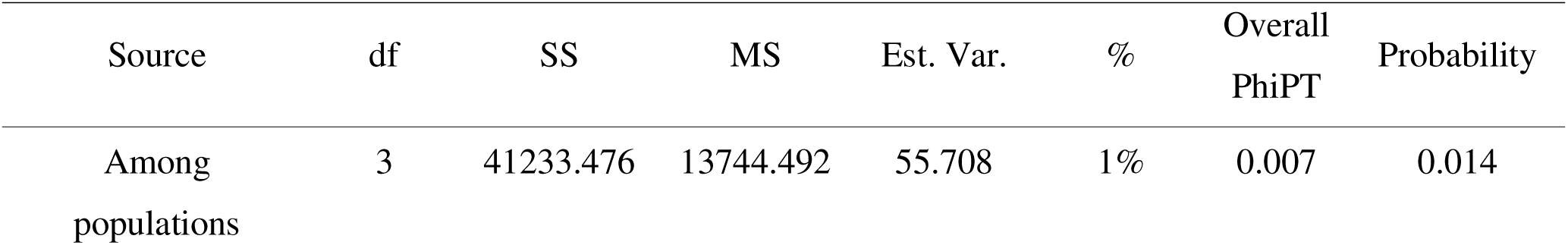
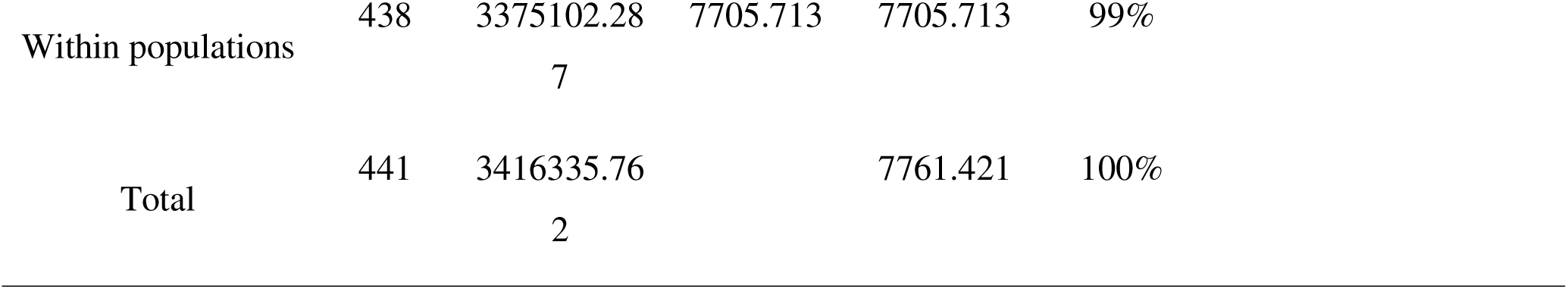
Analysis of molecular variance (AMOVA) of *Fusarium graminearum* residue populations according to sampling stages collected in four wheat fields in Brittany, France.

### Trichothecene and zearalenone chemotype profiling of *Fusarium graminearum*

Of the 335 CCs tested, 331 (98.8%) had the 15ADON chemotype, while 4 (1.2%) had the NIV chemotype. No isolates of the 3ADON chemotype were detected. The NIV chemotype originated from wheat grains and maize residues of F3 in Y1 and residues of F4 (both Y1 and Y2). Additionally, the ZEA chemotype was found for 288 CCs (86.0%). We subsequently assessed the degree of genetic differentiation between chemotype groups (15ADON vs. NIV and ZEA-positive vs. ZEA-negative) using an AMOVA analysis. For ZEA, the majority of genotypic diversity (99%) was observed within chemotype groups, with only 1% of the variation occurring among groups, and a slightly significant PhipT value was obtained (0.013, *p* < 0.05). No significant genetic differentiation was observed between different groups based on trichothecene chemotypes.

## Discussion

The aim of this study was to evaluate the genetic and chemotypic diversity within *F. graminearum* populations across different geographical regions and substrates (wheat grains and maize residues) and investigate the role of maize residues in contributing to the infection of wheat heads.

Overall, our results revealed high levels of genetic diversity without genetic differentiation according to field location and substrate type. This observation aligns well with previous studies reporting high genetic variability in *F. graminearum* across different geographic scales, from single fields to entire countries (Aamot et al. 2015; Burlakoti et al. 2017; Karugia et al. 2009; Kelly et al. 2015; Oghenekaro et al. 2021; Talas and McDonald 2015). In our case, a high level of genotype diversity was observed at the field scale in both wheat heads and maize residues (with a percentage of unique MLG per field ranging from 78 to 90%), indicating that a wide range of *F. graminearum* strains colonized these substrates over a cropping period.

The low differentiation between *F. graminearum* populations, despite geographical distances, as well as high gene flow between populations, indicates that these populations likely belong to one metapopulation. Similar lack of geographic genetic differentiation was observed in studies of *F. graminearum* populations from wheat and corn fields in China (Zhang et al. 2012), Norway (Aamot et al. 2015), Germany (Talas et al. 2011), and Canada (Burlakoti et al. 2017; Oghenekaro et al. 2021). Long-distance dissemination of inoculum through biotic (e.g., insects) and abiotic factors (e.g., wind) (Momeni and Nazari 2016), along with plant material transport, likely contributes to the absence of geographic genetic structure and high diversity in *F. graminearum* populations.

Sexual recombination is another key contributor to the large genetic diversity of *F. graminearum* populations within small field areas (Gale et al. 2011; Miedaner et al. 2001; Talas and McDonald 2015). Sexual reproduction is a critical part of the life cycle of *F. graminearum* (Trail 2009). An outcrossing rate of 35% was reported by Bowden and Leslie (1999) under laboratory conditions. Sexual reproduction primarily occurs on infected crop residues during the wheat flowering stage, generating a large amount of inoculum that subsequently infects wheat heads (Trail 2009). However, in the present study, the LD indices did not support the hypothesis of random mating in populations based on fields and sampling stages. The observed LD in the *F. graminearum* populations from wheat heads and maize residues from these fields suggests that sexual recombination has not occurred at a sufficient level for these populations to be considered as randomly mating. Nonetheless, we cannot exclude the hypothesis of a combination of both asexual and sexual modes of reproduction within *F. graminearum* populations.

In addition to sexual recombination, other various processes, such as mutations, gene flow, gene drift, population mixing, and asexual recombination through anastomosis, could also contribute to the observed high level of genetic diversity in the *F. graminearum* population (Talas and McDonald 2015). High gene flow between *F. graminearum* populations was previously reported as a dominant evolutionary force shaping the metapopulation (Talas and McDonald 2015; Yerkovich et al. 2020). Wind-dispersed ascospores produced through sexual reproduction could further support high gene flow between different fields, preventing population sub-structuring. Ascospores can travel long flight distances because of their structural characteristics; in contrast, macroconidia, which are fusiform in shape and formed in slimy masses, are typically transported over shorter distances, primarily by rain-splash and, to a lesser extent, by wind (Trail 2009). The supposedly low rate of sex among the studied populations also supports the hypothesis of a larger, interconnected population.

Furthermore, changes in biotic and abiotic pressures, such as fungicide application, crop rotation, the introduction of novel resistant cultivars, or variations in temperature, rainfall, and humidity, could also alter the current population structure by introducing new alleles, thereby shifting allelic frequencies and contributing to high genetic diversity within population (Talas and McDonald 2015). The adaptability of *F. graminearum* populations could be enhanced by the influx of new alleles, as multiple phenotypic traits, including pathogenicity, aggressiveness, and saprophytic behavior, may vary depending on the previously infected host (Akinsanmi et al. 2007). Balancing selection between the saprophytic (notably in residues) and parasitic phases (notably during wheat head colonization) of the pathogen’s life cycle, combined with low host selection pressure, allows a diverse range of isolates to survive and coexist (Miedaner et al. 2001, 2008). Therefore, the genetic diversity of *F. graminearum* populations is not exclusively a result of asexual reproduction, but is influenced by multiple factors, all of which support high genetic variation within these populations.

Phenotypic traits of *F. graminearum*, here trichothecene chemotype, also appear unaffected by the geographic locations of isolates. Various studies have documented a geographical partitioning of trichothecene chemotypes throughout Europe, with 3ADON dominating in Northern Europe (Aamot et al. 2015; Langseth et al. 2001; Nielsen et al. 2012; Yli-Mattila et al. 2009), whereas 15ADON appears predominant in Western and Southern Europe (Audenaert et al. 2009; Boutigny et al. 2014; Jennings et al. 2004; Pasquali et al. 2010; Prodi et al. 2009; Somma et al. 2014; Stępień et al. 2008; Talas et al. 2011). Accordingly, in this study, 99% of *F. graminearum* isolates from wheat grains and maize residues were of 15ADON type, indicating that this chemotype remains the almost exclusive one, whatever the substrate type or sampling stage in Brittany, France. Although NIV-producing isolates have been previously reported on wheat in European countries at various frequencies, ranging from 1 to 25%, only six *F. graminearum* isolates in the present study were predicted to produce NIV. The distribution of trichothecene types across Europe is potentially influenced directly or indirectly by multiple factors, including climate (Belizán et al. 2019; Madgwick et al. 2011), regional host composition (Burlakoti et al. 2017; Nielsen et al. 2012), agricultural practices (Crippin et al. 2020), and the historical or recent movement of pathogen populations into new areas (Oghenekaro et al. 2021; Talas and McDonald 2015; Ward et al. 2008). Despite that we found no evidence of the presence of 3ADON chemotypes in Brittany, the shift from 15ADON to 3ADON, which is more aggressive and toxigenic, has been observed in the USA (Gale et al. 2007; Puri et al. 2016; Ward et al. 2008) and Canada (Ward et al. 2008). This underscores the need for regular temporal and spatial surveys of trichothecene chemotypes to optimize regional FHB management strategies.

This study also aimed to determine the transfer of inoculum from residues to wheat heads. The absence of substrate genetic differentiation and high genetic similarity between *F. graminearum* residue and grain populations, irrespective of geographical locations, supports the fact that residues contributed to FHB infection. This finding corroborates with previous studies showing the tillage effect on FHB incidence and DON contamination in cereal grains (Blandino et al. 2010; Drakopoulos et al. 2021). However, we failed to demonstrate any field-specific dispersal pattern from residues to grain, most probably because our *F. graminearum* populations belong to a metapopulation at least within our sampled area, or due to insufficient sampling depth. This unexpected finding may indicate a significant inoculum exchange between fields, suggesting that long-distance transfer contributes to the dissemination of *F. graminearum*. Our results showed that the inoculum of *F. graminearum* was able to move over large distances (although the time span is unknown), as previously demonstrated (Gale et al. 2002; Naef and Défago 2006; Zeller et al. 2004). These authors identified large, interbreeding populations of *F. graminearum* with little to no subdivision in the USA, Europe, and Eastern China. Additionally, high genetic similarity was detected in *F. graminearum* populations collected 11 years apart from maize residues and wheat, despite being separated by a geographical distance of 100 km (Naef and Défago 2006). Even though we provided evidence that residue-borne inoculum likely contributed to FHB incidence in our fields, we cannot rule out the possibility of airborne inoculum from neighboring fields contributing to FHB. Notably, since the F1 field was plowed and residues were left on the soil surface, this underlines that F1 grain-associated populations originated from airborne inoculum from short and/or long-distance dispersal events.

## Conclusion

In conclusion, this study revealed significantly high levels of genetic diversity within *F. graminearum* populations from wheat grains and maize residues, with low genetic differentiation and high gene flow across geographic locations in Brittany. These findings suggest that *F. graminearum* populations likely form a single metapopulation, reinforcing the importance of regional gene flow in shaping population structure. Notably, the high genetic similarity between populations from wheat grains and maize residues underscores the critical role of maize residues as a significant source of inoculum for wheat head infection, regardless of geographic origin. This result highlights the need to consider both local and distant sources of inoculum in developing more effective FHB management strategies. Spatio-temporal dynamics studies conducted over an extended period of time are needed to infer the distance dispersal of *F. graminearum* more clearly. In addition to the significant contribution of crop residues, it is important to explore the potential contributions of other local inoculum sources of *F. graminearum*, such as wild grasses or weeds. Furthermore, the use of spore traps should also help determine the contribution of airborne inoculum. Understanding the contribution of these factors to the pathogen’s population structure will be crucial for refining disease management approaches.

## Acknowledgements

We thank the farmers for their participation and for kindly giving us access to their fields to take samples.

## Data availability

**Figure S1.** Multilocus genotypes (MLGs) of twelve *Fusarium graminearum* populations from four different wheat fields and two types of substrates (wheat grains and maize residues) across two wheat cycles in Brittany, France, based on sequence-based microsatellite genotyping of 34 markers. Populations are denoted by color, and the y-axis shows the counts of isolates for each MLG in the total dataset.

**Figure S2.** Clonal complexes (CCs) of twelve *Fusarium graminearum* populations from four different wheat fields and two types of substrates (wheat grains and maize residues) in Brittany, France. The genetic population dataset was obtained based on sequence-based microsatellite genotyping (SSRseq) of 34 markers. The CCs of *F. graminearum* populations were identified at the triple locus variant (TLV) level using the goeBurst algorithm implemented in the Phyloviz version 2.0 software. Populations are denoted by colors, and the y-axis shows the counts of isolates for each CC in the dataset.

