## Supplementary figures and images for "A population genetic study of *Fusarium graminearum*, the causal agent of Fusarium Head Blight, provides insights into patterns of inoculum dispersal in wheat fields at a regional scale"

### Supplemental Figure S1

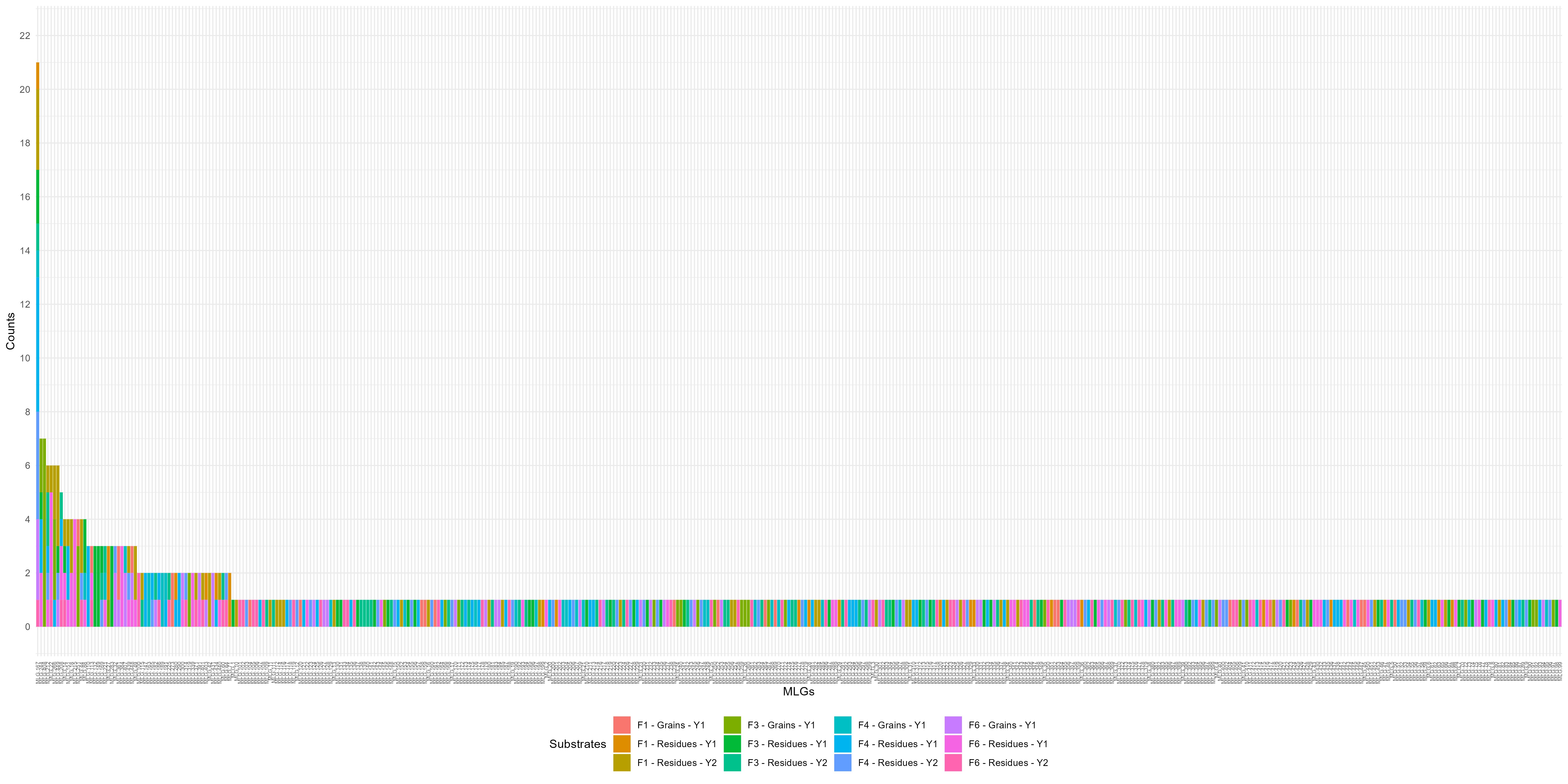

### Supplemental Figure S2

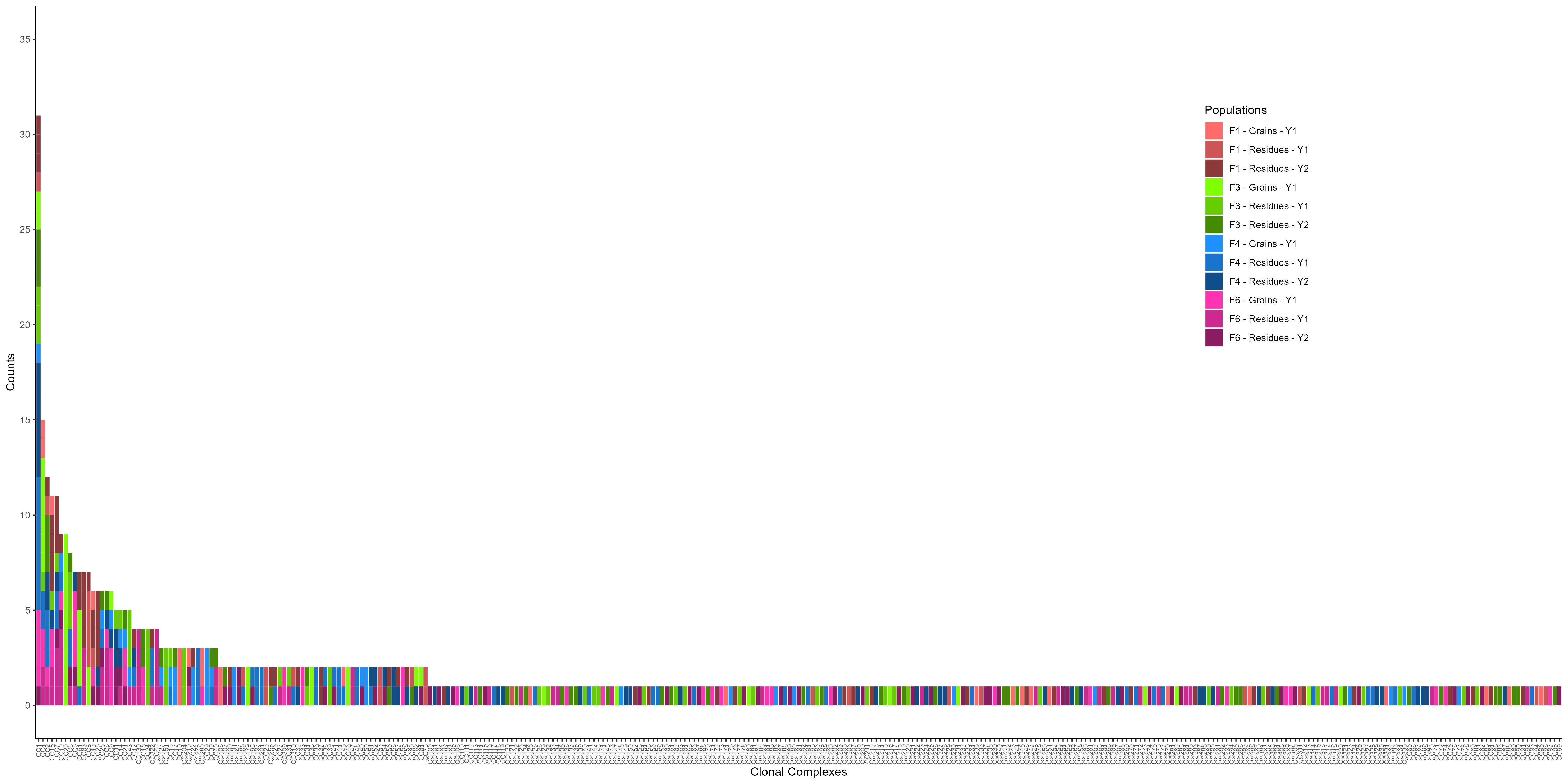
